# Path tortuosity changes the transport cost paradigm in terrestrial animals

**DOI:** 10.1101/2020.08.20.259390

**Authors:** R. P. Wilson, K. A. R. Rose, R. S. Metcalfe, M. D. Holton, J. Redcliffe, R. Gunner, L. Börger, A. Loison, J. Miloš, M. S. Painter, V. Silovský, N. Marks, M. Garel, C. Toigo, P. Marchand, N. C. Bennett, M. A. McNarry, K. A. Mackintosh, M. R. Brown, D. M. Scantlebury

## Abstract

Animal movement paths are variously tortuous, with high turn rates predicted to be energetically costly, especially at high speeds. Animals travel most efficiently at the speed that gives the lowest cost of transport (COT), a well-defined point for movement in fluid media. However, theoretically, land animals should travel at their maximum speed to minimize COT, which they do not, instead travelling at walking pace. We measured oxygen consumption in humans to demonstrate that the energetic costs of turning increase disproportionately with both speed and angular velocity. This resulted in the minimum COT speed occurring at very low speeds, which reduced with increased path tortuosity. Data on turn rates from six free-ranging terrestrial species underpinned this because all individuals turned faster at the slowest speeds across the full speed range. The optimum movement speed for minimum COT in land animals thus depends on the environment and behavior since both affect track tortuosity.

## Introduction

Reaction to stimuli by movement is a fundamental tenet of the animal kingdom, with appropriate responses enhancing survival and lifetime reproductive success [1]. Movement requires energy, which varies according to the speed of the animal [2]. The speed that an animal selects for movement, and therefore the energy designated to it over time, depends on the purpose of the movement [3]. There are three primary reasons for this: (i) animals may move at any speed, including their maximum, to evade predators or capture prey [4], to maximize net rate of acquisition of food energy [5], or to provide young with food at an appropriate rate for their growth [6]. (ii) animals may move at a speed that minimizes power costs (a concept that is primarily applicable for flying animals) [7]. In addition, (iii) animals may move at a speed that minimizes the cost of transport (COT_min_), maximizing the distance travelled per unit of energy [8, 9]. This last paradigm [10] is particularly important because most travel is assumed to be under strong selection pressure for animals to maximize output while minimizing input, which occurs at COT_min_ [e.g. 11, 12]. COT_min_ is well defined for any instance where power for movement increases with speed at a greater than linear rate, such as in aquatic and volant animals. Indeed, wild animals in fluid media generally travel at these speeds [13]. However, beyond basal metabolic rate [14], most terrestrial animals have a linear relationship between power and speed [10, 15] and therefore, because COT is derived by dividing the power by the speed, COT in these animals will always decrease with increasing speed. In short, the lowest theoretical costs of transport in terrestrial animals should occur at their maximum speeds (but see [16, 17]). Yet these are not the speeds at which animals normally travel in natural contexts [18]. Although the probability of accident and injury increases at very high speeds (e.g. [19]), which is a clear reason to avoid them unless there is a compelling motive to sprint (e.g. in cursorial predators and their prey [20]), there is currently no energy-based explanation as to why land animals travel through their environments at their normal chosen speeds.

Based on recent studies that found that the costs associated with turning are substantial [21, 22], we hypothesized that track tortuosity should therefore alter the speed of COT_min_ in terrestrial animals. To test this, we set up experiments to gain empirical data on how speed and angular velocity in humans relate to energy expenditure and thereby to the cost of transport. At the same time, we equipped 6 species of free-living animal with tags that allowed us to study animal speed in tandem with angular velocity to see whether our physiological findings translated into patterns of movement in the natural environment.

## Results

### Empirical data from humans show how increasing track tortuosity reduces the minimum cost of transport speed to a slow walk

We examined the energetics of moving in humans by measuring oxygen consumption in 20 subjects walking defined trajectories consisting of varying angles between straight-line sections at defined speeds. We noted how power increased disproportionately with increasing angular velocity, which resulted in clear cost of transport minima that did not occur at the higher walking speeds. Rather, minimum costs of transport occurred at lower walking speeds when angular velocities were high.

Our calculations of power-use by participants walking variously angled courses showed a significant interaction effect between walking speed and turn angle on total mass-specific metabolic power (speed^2^ × angle category, F = 101.81, P<0.001), with the rate of increase in power for a given increase in travel speed increasing disproportionately with turn angle (Fig 1a, Table 1). Consequently, there was a significant interaction between walking speed and turn angle on the total mass-specific cost of transport (speed^2^ × angle category, F = 59.13, P<0.001) (Fig. 1b, Table 1). The cost of transport curve showed distinct minima for paths incorporating turns, becoming more U-shaped, and with COT_min_ shifting to progressively lower walking speeds with increasing turn angle (Fig 1b).

**Table 1.**
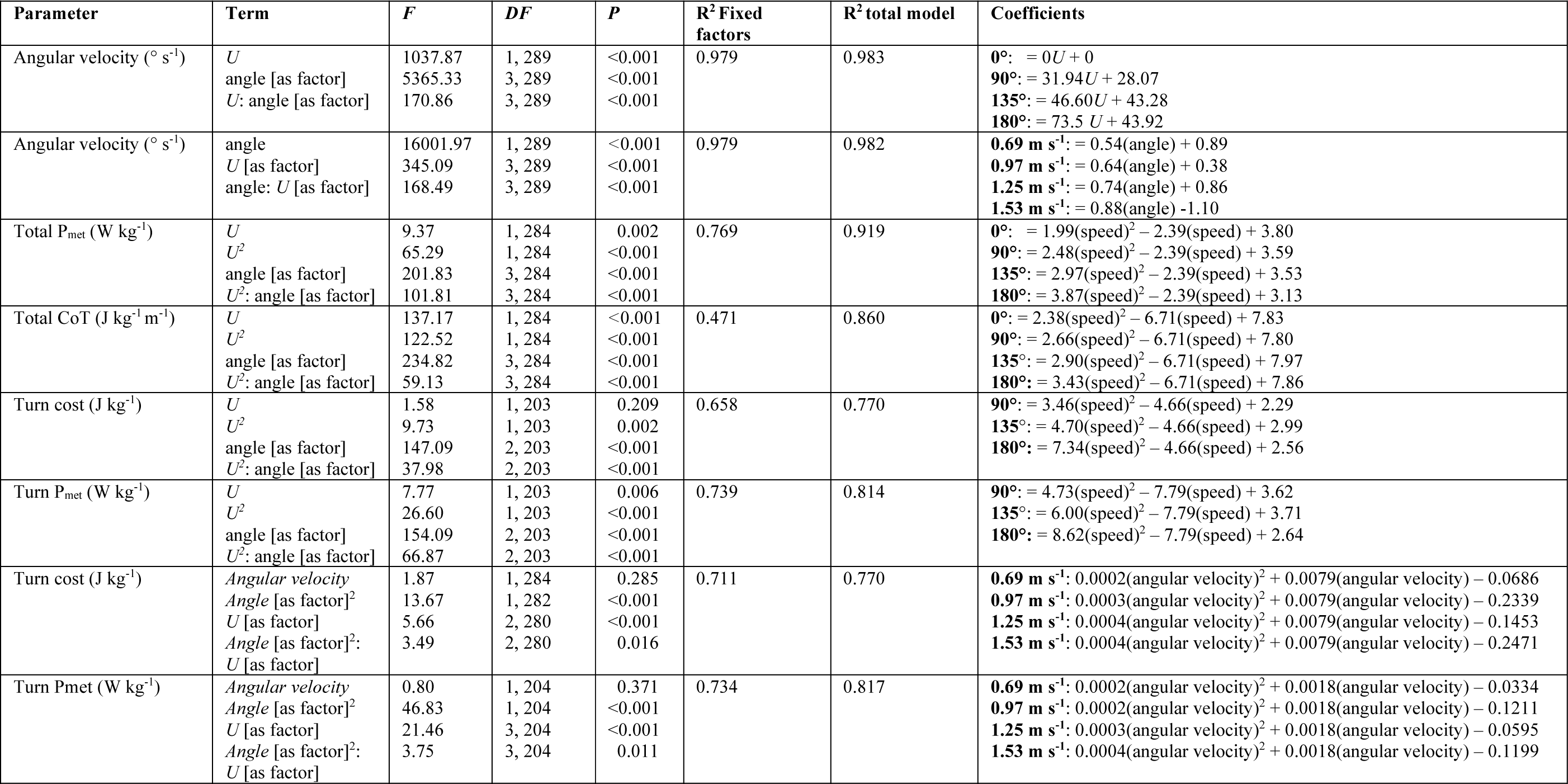
Summary of the outputs of linear mixed-effects models conducted to investigate the effects of speed and angle on angular velocity or metabolic parameters and the effects of speed and angular velocity on metabolic parameters.

**Table 2.** Details of tag deployments on free living animals

| Species | No. | Study area | Tag weight (g) | Tag weight as % body weight | Deployment period | Deployment length (days) | Ethical permissions |
| --- | --- | --- | --- | --- | --- | --- | --- |
| *Panthera leo* | 5 | Kalahari, S. Africa | 1240-1330 | <2 | Feb-19 | 18-21 | Queen's University Belfast (QUB-BS-AREC-18-006), Pretoria University (NAS061-19), South African Parks (011-18) |
| *Rupicapra rupicapra* | 5 | Les Bauges, France | 450 | 1.13 | June 2017-May 2018 | >250 days | IP 1516-14 |
| *Capra ibex* | 5 | Belledonne, France | 450 | 0.64 | April 2017-March 2018 | >250 days | IP 1516-14 |
| *Ovis gmelini* | 4 | Caroux-Espinouse, France | 450 | 0.90 | May 2017-April 2018 | >250 days | IP 1516-14 |
| *Sus scrofa* | 5 | Kostelec, Czech republic | 850 | 0.8-1.3 | May - Oct 2019 | 30-120 | Ministry of environment CZ (MZP/2019/630/361) |
| *Capra aegagrus* | 5 | Les Bauges, France | 510 | 1.28 | August-September 2017 | 29 days | SU-Ethics-Student-190717/246 |
| *Homo sapiens* | 20 | Swansea, UK | <1 | <1 | July-November 2019 | < 1 day | Swansea University CoE ethics: 2012:075 |

**Fig 1.**
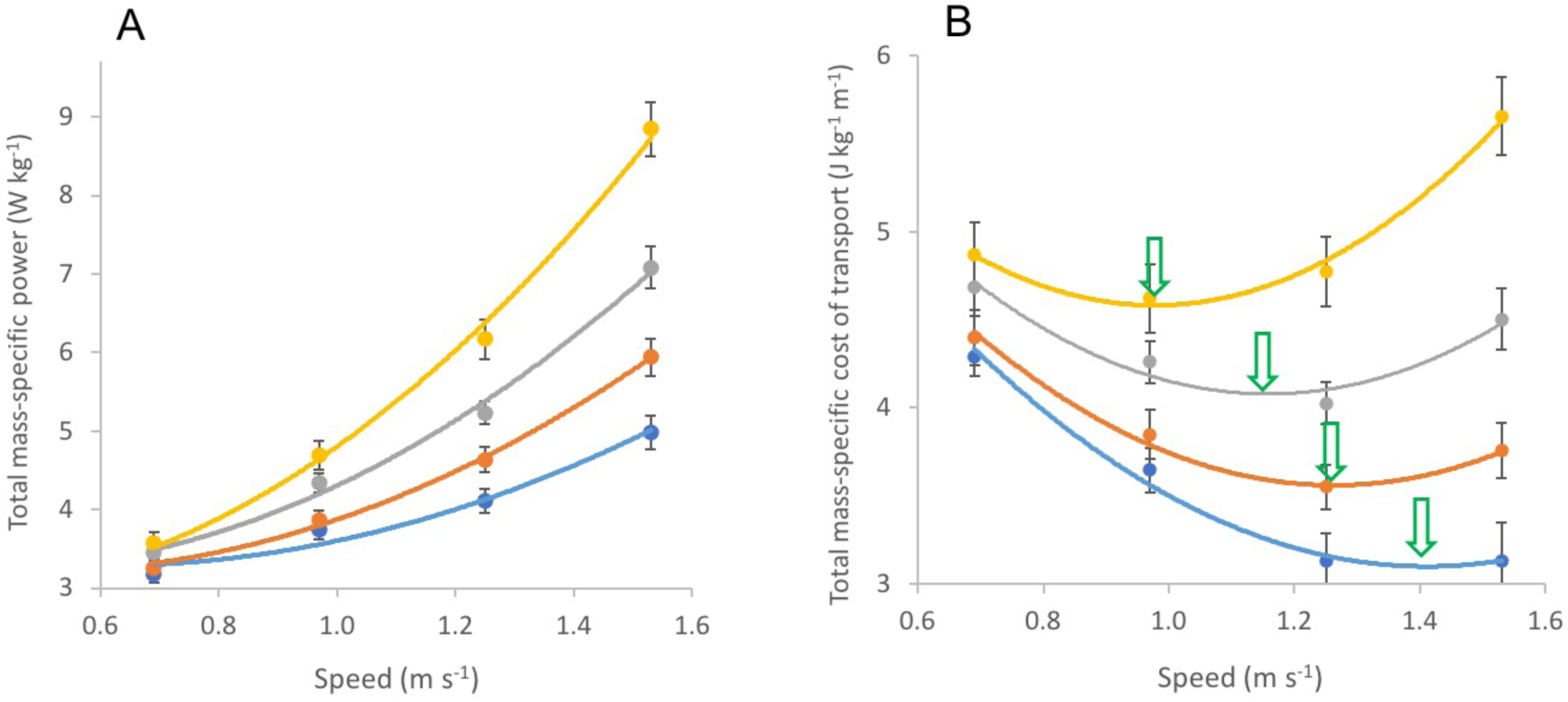
Metabolic measurements *versus* average walking speed for 20 human participants walking tracks with turns of varied extent. (A) Mean (± SE) total mass-specific power and (B) mean (± SE) total mass-specific cost of transport for straight movement (blue), 90° turns every 5 m (orange), 135° turns every 5 m (grey) and a 180° turns every 5 m (yellow). Quadratic curves were fitted using the coefficients of the outputs of linear mixed-effects models (see Table 1). Arrows show COT_min_.

Similarly, significant interaction terms were found for the quadratic relationships between speed and turn cost (J kg^-1^, speed^2^ × angle category, F = 37.98, P<0.001) and turn power (W kg-1, speed2 × angle category, F = 66.87, P<0.001) (Table 1), with the rate of increase with speed becoming greater as angle increased (Fig 2).

**Fig 2.**
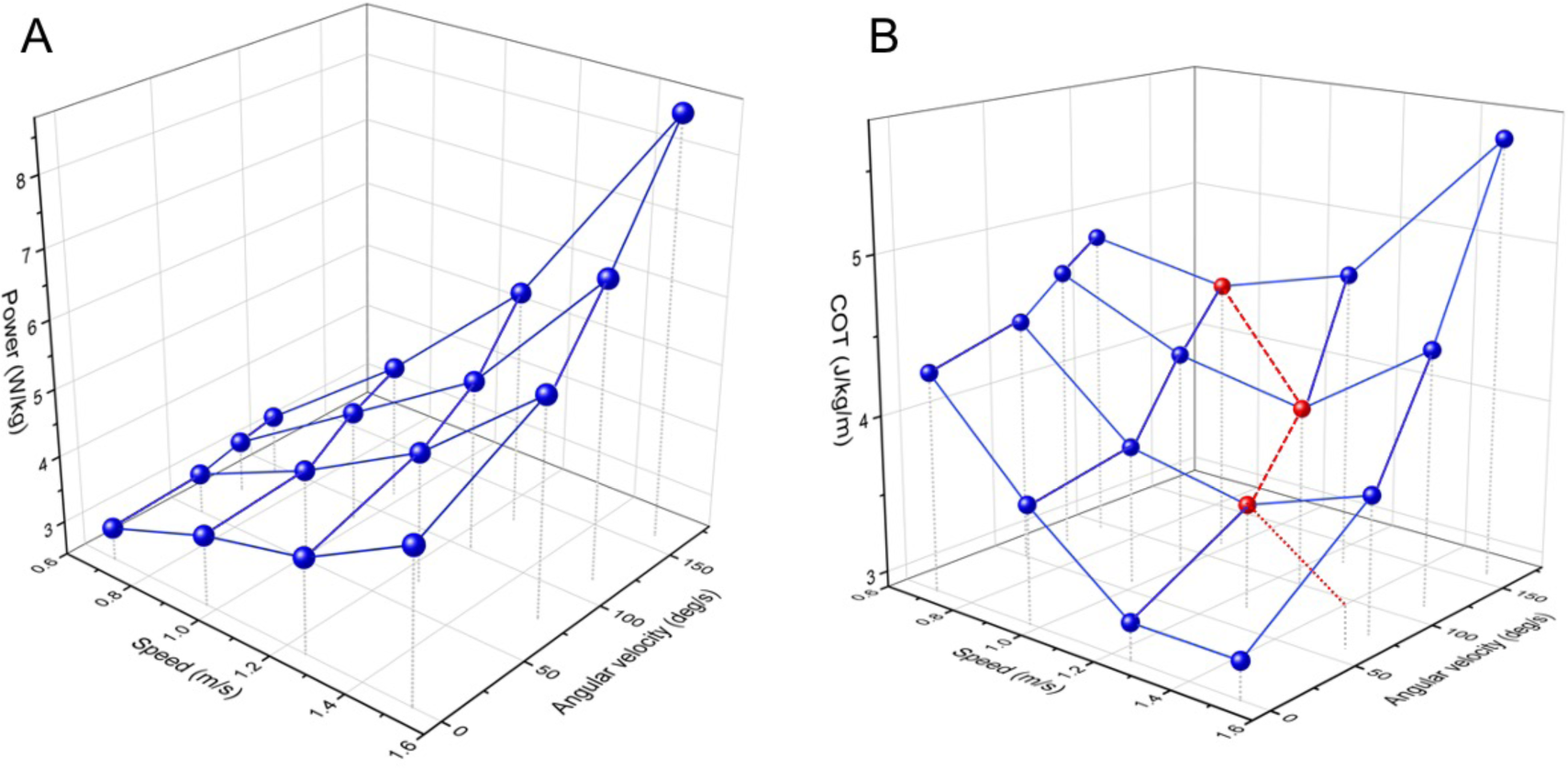
3D representations of how mass-specific metabolic power and the mass-specific cost of transport are influenced by average travel speed and angular velocity. (A) Mass-specific power, and (B) mass-specific cost of transport (COT) for twenty male participants walking courses with defined angle turns interspaced with straight sections. Each point represents a mean value from twenty values, each of which is derived from a single participant. Power costs increase with both speed and angular velocity. Against this, the accelerating power costs of higher angular velocities produce cost of transit minima (highlighted in red) where any turning occurs and these minima occur at lower speeds with higher angular velocities.

### Wild animal data support the idea that normal travel speed varies with angular velocity

We fitted five wild mammal species (African lions *Pathera leo*, chamois *Capra rubicapra*, ibex *Capra ibex*, mouflon *Ovis orientalis*, wild boar *Sus scrofa*) and one domestic animal (the goat *Capra aegagrus*) allowed to range freely, with devices that enabled us to record speed (or a proxy for it) and angular velocity. All individuals tended to travel relatively slowly, but they predominantly engaged in the highest angular velocities at the lowest speeds (using dynamic body acceleration as a proxy for speed [23]) decreasing angular velocities in their turns as their travel speeds increased (Fig 3). The incidence of turning behavior was apparent in examples of the free-living animal movement data when resolved at sub-second level, which showed intensive and extensive turning behavior, even when movement paths appeared directed at larger scales (e.g. Fig 4).

**Fig 3.**
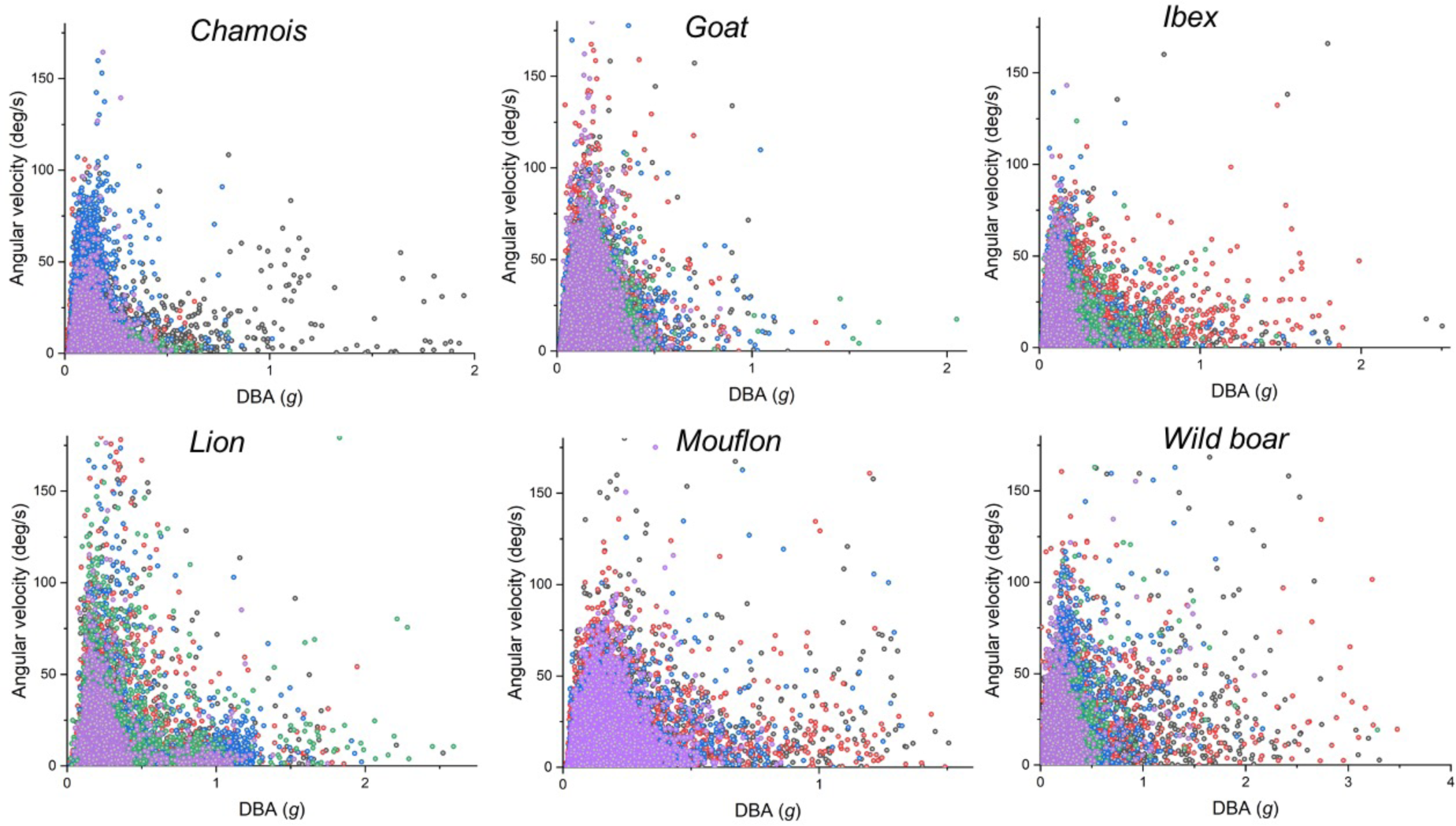
Incidence of angular velocity against dynamic body acceleration (DBA) – a linear proxy for travel speed – in 6 free-living terrestrial species. The lion data show both DBA [black numbering on x-axis] and speed [red numbering] because these animals were equipped with high resolution GPS – see inset). Each point shows a value taken at *ca.* 10 s intervals over a length of time that corresponds to about 20,000 points for each individual (N = 5 per species except for the mouflon where N = 4, with individuals depicted by different colors). Note how all distributions show decreasing angular velocities with increasing speed across all speed ranges, starting with the slowest.

**Fig 4.**
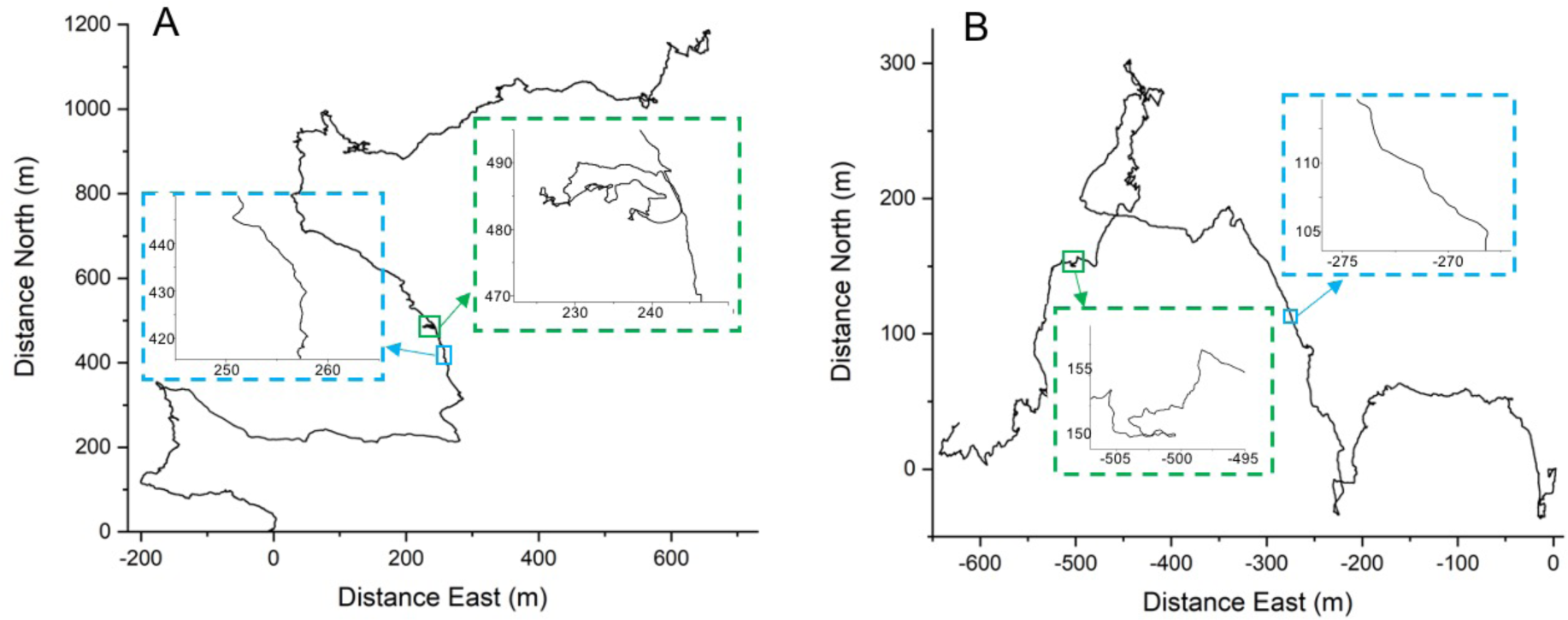
Examples of 20 h movement paths (at 4 Hz and elucidated by dead-reckoning) of two of the study animals. (A) a chamois and (B) an Ibex, showing in the insets the scale over which turns are made (cf. our VO2 study on humans). Note how even apparently straight-line paths (blue boxes) show clear turns, although angles are obtuse, while tortuous sections (green boxes) show both obtuse and acute turns.

## Discussion

There have been extensive studies of the relationship between terrestrial animal speed and power, the vast majority of which have been conducted on a treadmill to show that power increases linearly with speed [10]. While fundamental, this highly controlled scenario does not echo the situation found in nature where variability in, for example, topography and surface penetrability will also affect power to travel [24] and thereby the cost of transport. Interestingly, in a recent study of how scaling affects costs in animals moving on inclines, Halsey and White [25] present data that show clear minima in costs of transport with slope (although the effect of speed is not detailed). This, therefore, already points to an important effect of how the environment is expected to structure selection of animal speed for energetically optimum movement. But animals turn frequently, as our Figure 4 shows, and presumably much more than they encounter varying inclines. Previous works have indicated that turns are energetically costly [21, 22] but, due to their experimental protocol, these could not have defined how the costs of transport relate to angular velocity and speed, nor that angular velocity could have created clear minima in costs of transport at such low speeds. The extraordinary magnitude of the increase in power costs with increasing angular velocity and speed is, in part, explained by animals having to develop forces to counteract the centripetal force incurred in the turn, given by; F = m.v^2^/r, where m is the mass, v, the velocity and r the radius of the turn. Thus, some of the energy used for turning has to be used in developing lateral forces. However, acute turns will also require some forces opposing the drive force, being manifest as deceleration followed by corrective acceleration for straight-line travel after a turn. This will occur even at low speeds, which explains the sharp drop off in angular velocity in turns, even at low speeds in our wild animals (Fig 3) and the minimum cost of transport occurring at such low walking speeds of humans during 180° turns (Fig 1).

The high power associated with turns translates into well-defined minima in the costs of transport for defined speeds and angular velocities (Fig 2b). This means that an overall optimum speed for a wild animal, moving to minimize transport costs, will depend on the frequency and extent of turns. These will depend on a large number of turn elicitors that confer benefit to the animal if it turns. These will range from barriers, including localized ones such as trees/shrubs (e.g. [18]), or navigating through differential energy landscapes [24], through to acquisition strategies for food, such as area-restricted search [26], or mates [27]. Indeed, we suggest that the incidence of turning behavior in wild animals has been underestimated, not least because we have little capacity to resolve it at the scale necessary to be energetically meaningful, even using GPS technology [28]. A notable exception is with inertial/dead-reckoned data (e.g. [29]), which show that animal paths can consist of extensive and intensive turning behavior (e.g. Fig 4).

Variation in the environment or in animal behavior would also lead us to expect animals to vary travel speed accordingly, even intra-specifically, reflecting different motivations for speed selection [3], which is consistent with the animal data we present here (Fig 3). Clearly, although such data may not always agree with animals actually using COT_min_ for a number of reasons [3], the interplay of speed and angular velocity, and the huge effect that they have on movement energetics, points to the importance of both in modulating trajectories according to function. Indeed, future treatises that attempt to link this with the multiple other factors that affect the energy costs of travel (the energy landscape [24] may find that we can explain animal speed- and turn velocity-selection much more satisfactorily than we can at present.

Our treatment is simplistic for a number of reasons. Not least, we only consider power and COT metrics as bi-dimensional when they will be affected by multiple properties of the environment that modulate power use, such as slope and surface penetrability [24]. This will be further complicated by gait changes [30–34]. This work nevertheless demonstrates the profound effect that turns have on cost of transport for terrestrial animals using humans as a model, although we expect the principle to be the same for flying and aquatic animals. If we are to progress with models purporting to help us understand animal trajectories from an energetic, and ultimately behavioral, perspective though, we will have to accept that small things, such as acute corners can sometimes make big differences in understanding the details and elicitors of movement trajectories [35].

## Materials and Methods

### Participants

Twenty able-bodied male participants (mean ± SD: age 30 ± 8 years; body mass 75.6 ± 11.2 kg; stature 1.78 ± 0.06 m; body mass index: 24.0 ± 3.2 kg·m^-2^) gave their written informed consent to volunteer for this study. Ethical approval was granted by the Swansea University ethics committee. Stature (Holtain, Crymych, UK) was measured to the nearest 0.01 m and body mass (Seca, Germany) to the nearest 0.01 kg.

### Trials

Participants walked at four different speeds (0.69, 0.97, 1.25 and 1.53 m s^-1^) and four turn angle extents (0, 90, 135 and 180°), giving a total of sixteen speed and angle combinations. Separate walking tracks were marked onto a flat-surfaced sports lab floor using tape for each angle of interest. Each track alternated between 5 m straight lines and turns with equal numbers of left and right turns (0°: a 25 m straight line with markers every 5 m; 90°: two squares sharing one 5 m straight; 135°: two eight-angled star polygons sharing a 5 m straight; 180°: a 5 m straight). Participants’ average walking speed was controlled using a metronome at either 8, 12, 15 or 18 bpm. Participants walked barefooted, carried out only step-turns (not spin-turns), and reached a corner (or 5 m mark) on each metronome beat (i.e. travelled 5 m between beats). Walking conditions were block-randomized by turning angle: all speed trials for a given angle were conducted before moving on to the next angle, but average travel speed and the sequence in which the angle conditions were completed by each participant were randomized and counterbalanced. Each walking condition lasted 3 minutes to allow achievement of a metabolic steady state, and each condition was separated with 3-minutes of quiet seated rest.

### Respirometry

A portable and wireless breath-by-breath respirometry system (MetaMax 3B, Cortex, Biophysik, Leipzig, Germany) was used to measure rates of oxygen consumption (VO2, L•min-1•kg-1) and carbon dioxide production (VCO_2_, L·min^-1^·kg^-1^). Prior to each trial, reference gases were used to calibrate the O_2_ and CO_2_ analysers and the turbine volume transducer was calibrated using a 3 L syringe (Hans Rudolph, Kansas City, MO). Respiratory exchange ratios (VCO_2_:VO_2_) and their thermal equivalents taken from (36) were used to convert VO_2_ to metabolic power (*P*_*met*_, W·kg^-1^). The cost of transport (J·kg-1·m^-1^) was calculated by dividing *P*_*met*_ by average travel speed (m s^-1^).

### Identifying angular velocity from turns

All participants were equipped with a lower back, mid-line mounted ‘Daily Diary’ tag (37) recording tri-axial (orthogonal) acceleration and magnetic field intensity at 40 Hz. Post-experiment, all magnetic field data were plotted on a tri-axial spherical plot of magnetic field intensity (m-sphere; see (38) for details). While participants walking in straight-lines led to little variation in tri-axial plotted position on the m-sphere, those that executed turns resulted in rapid movement of points across the m-sphere surface which clearly showed at which points turns began and ended so that turn duration could be derived. Maximum resolution for this was 1/40^th^ second. The duration of turns was determined for 5 turns per individual per speed and turn angle combination.

### Derivation of costs

Metabolic measurements for the 0° trials were corrected for the costs of 180° turns that were necessary at the end of every 25 m of straight-line walking. In order to calculate the cost of transport for straight line walking, a series of calculations were undertaken. Firstly, the time *t*, to travel one length (either 5 m or 25 m) and execute one turn for any given track was calculated as distance travelled divided by speed, 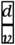. The total mass-specific cost of walking one length and executing one turn was then calculated as the time taken × Pmet (*tP*_*met*_). The total mass-specific cost of walking in a straight line for 20 m was then calculated by subtracting the mass-specific cost of walking 5 m + 1 turn in 180° trials from 25 m + 1 turn in the 0° trials; *T*_*CoW*_ = *tP*_*met*_|_*d*=25_ − *tP*_*met*_|_*d*=5_. The total mass-specific cost of walking in a straight line for 20 m was then divided by 20 in order to derive the total mass-specific cost of transport (J kg^-1^ m^-1^). Turn costs for any given speed and angle combination were calculated by subtracting 5× the cost of transport for straight-line walking from the total mass-specific cost of walking 5 m and executing one turn. Each participant’s turn costs were divided by their mean turn durations in order to calculate turn power (W kg^-1^).

### Animal tagging

Five wild mammal species (African lions *Pathera leo*, chamois *Rupicapra rupicapra*, ibex *Capra ibex*, mouflon *Ovis orientalis* and wild boar *Sus scrofa*) and one domestic mammal (domestic goat *Capra aegagrus*), which was allowed to range freely, were fitted with collar-mounted GPS-enabled ‘Daily Diary’ tags (*2*) recording tri-axial magnetic field intensity and tri-axial acceleration at 10-40 Hz (see species-specific details in table 1). Tags were left in place for between 15 and >250 days before being retrieved. Downloaded data were treated to determine movement patterns using dead-reckoning as described in (29), correcting for drift using the periodic GPS positions (29). From these data, a random period (24-50h) was isolated from each individual, with some variation in durations to ensure that appreciable movement had taken place. For these periods, the vectorial dynamic body acceleration (VeDBA), a good proxy for speed (23, 39), was calculated according to (40) as well as the angular velocity over one second at approximately 10 s intervals so as to give ca. 20,000 angular velocity-VeDBA pairs per individual. Data from five individuals for all species were used in this manner except for the mouflon, where only 4 animals were used.

### Statistical analyses

Linear mixed-effects models (lme4 package in R Studio v 1.2.1335) were conducted in order to investigate the relationship between angular velocity and average travel speed with differing angle extent; angular velocity with angle extent at different speeds, mass-specific metabolic parameters and speed with different angle extent; and mass-specific metabolic parameters with angular velocity at different speeds or angle extent. Participant identity was included in the models as a random factor. The F statistic and marginal and conditional R^2^ were determined using the ‘car (3.0-5)’ and ‘MuMIn (1.43.6)’ packages, respectively.

## Acknowledgements

RPW conceived the overall study while KAR, RSM developed and executed the human-based work with support from MAM and KAM. DMS, NM, NCB, and RPW conceived specific ideas for the lion work and JR, RG, LB, AL, JM, MSP, VS, MG, CT, PM conceived methodologies and implemented various aspects of the fieldwork for the other species. DMS, NM and NCB acquired funding and support. KAR, RSM, JR, RG and RPW performed the analyses while MDH developed and provided software and analytical ideas. All authors contributed to the manuscript preparation.

## Notes

### Competing Interest Statement

The authors have declared no competing interest.

